# Local parasite pressures and host genotype may modulate epigenetic diversity in a mixed-mating fish

**DOI:** 10.1101/603274

**Authors:** Waldir M. Berbel-Filho, Carlos Garcia de Leaniz, Paloma Morán, Jo Cable, Sergio M. Q. Lima, Sofia Consuegra

## Abstract

Parasite-mediated selection is one of the main drivers of genetic variation in natural populations. The persistence of asexual reproduction and self-fertilization, however, challenges the notion that low genetic variation and inbreeding compromise the host’s ability to respond to pathogens. DNA methylation represents a potential mechanism for generating additional adaptive variation under low genetic diversity. We compared genetic diversity (microsatellites and AFLPs), variation in DNA methylation (MSAFLPs), and parasite loads in three populations of *Kryptolebias hermaphroditus*, a unique mixed-mating (partially self-fertilising) fish, to analyse the potential adaptive value of DNA methylation in relation to genetic diversity and parasite loads. We found strong genetic population structuring, as well as differences in parasite loads and methylation levels among sampling sites and selfing lineages. Globally, the interaction between parasites and inbreeding with selfing lineages influenced DNA methylation, but parasites seemed more important in determining methylation levels at the local scale.

## Introduction

Organisms with mixed-mating reproduction (alternating between self-fertilisation and outcrossing) can benefit from the advantages of both biparental and uniparental reproduction: outcrossing generates genetic variability and adaptability potential, while selfing ensures reproduction without partners (Jarne and Chalesworth 1993). Reproductive assurance (Darwin 1876) gives self-reproducing individuals an advantage when colonising new environments (Baker 1955) and ensures the genetic transmission of both sets of parental genes (Fisher 1941). The downside of selfing, however, is that the progeny can have reduced fitness compared to their outcrossed counterparts, and suffer from inbreeding depression (Charlesworth and Willis 2009). Thus, occasional outcrossing should be beneficial when inbreeding can impair offspring fitness (Damgaard et al. 1992; Maynard Smith 1978).

The Red Queen hypothesis (Van Valen 1973; Bell 1982) is often invoked to explain the occurrence of sexual reproduction in face of the advantages of asexual reproduction (Blirt and Bell 1987; Lively 1987; Lively and Morran 2014). According to this hypothesis, the more genetically diverse offspring of sexually reproducing individuals provide a “moving target” to parasites, making it more difficult for them to adapt compared to the “more static” offspring of asexual/uniparental individuals (Maynard Smith 1978; Hamilton 1980; Lively et al. 1990;). Yet, while sexual reproduction seems the general rule in animals (approximately 99%; Slowinski et al. 2016), asexual and self-fertilising lineages sometimes persist in a wide range of environments (Zhang et al. 2010), suggesting that their adaptation and long-term survival could be facilitated by other factors in addition to genetic variability (Verhoeven and Preite 2014).

Non-genetic factors (including epigenetic mechanisms) can play an important role in generating adaptive phenotypic variation (Bossdorf et al. 2008; Verhoeven et al. 2016; Bonduriansky and Day 2018), including resistance to parasites (Verhoeven et al. 2010; Wenzel and Piertney 2014). Epigenetic mechanisms (e.g. histone modifications, microRNAs, DNA acetylation), can modulate changes in gene expression in response to environmental variation without involving changes in DNA sequence (Bossdorf et al. 2008; Richards et al. 2017). DNA methylation is the best characterized epigenetic modification (Lea et al. 2017), and has important roles on pre-transcriptional control in several biological processes, such as cell differentiation and genomic imprinting (Koch et al. 2016). Variation in DNA methylation is not completely independent from the genome, and epialelles can have different degrees of autonomy from the genotype (Richards 2006; Dubin et al. 2015). In addition, in some plants and animals, individuals with low levels of heterozygosity display high levels of genome-wide DNA methylation variation (Richards et al. 2012; Schrey et al. 2012; Liebl et al. 2013), sugesting that DNA methylation could contribute to the adaptation of asexual and inbred organims with limited genetic diversity to environmental change (Castonguay and Angers 2012; Schrey et al. 2012; Liebl et al. 2013; Verhoeven and Preite 2014; Douhovnikoff and Dodd 2015).

Increasing evidence suggests that epigenetic mechanisms, including genome-wide DNA methylation, are involved in host-pathogen interactions (Gómez-Díaz et al. 2012; Hu et al. 2018), but the mechanisms are better known in plants than in animals (Annacondia et al. 2018; Hewezi et al. 2018; Gómez-Díaz et al. 2012). Mixed-mating organisms represent ideal models to test the associations between genetic and epigenetic variation with pathogen pressures because selfed and outcrossed offspring naturally coexist, usually displaying very different levels of genetic diversity. Negative associations between genetic diversity and parasite loads have been previously observed in mixed-mating animals (Lively and Morran 2014; Ellison et al. 2011), with inbred individuals usually harbouring more parasites. The relationship between epigenetic variation, parasites and mixed-mating, however, has not been explored.

Here, we compared genetic diversity, variation in DNA methylation, and parasite loads in three natural populations of the mixed-mating mangrove killifish *Kryptolebias hermaphroditus* distributed along the Brazilian coast (Tatarenkov et al. 2017). The genus *Kryptolebias* contains the only known mixed-mating vertebrate species (*K*. *marmoratus* and *K*. *hermaphroditus*), characterised by variable rates of selfing and outcrossing (Tatarenkov et al. 2017). Populations of both species consist mainly of self-fertilizing hermaphrodites and varying levels of males at low frequencies (Tatarenkov et al. 2017; Berbel-Filho et al. 2016), and exhibit high levels of homozygosity (Tatarenkov et al. 2009, 2017), suggesting that self-fertilization is the most common mode of reproduction (Avise and Tatarenkov 2015).

We analysed microsatellites (previously shown to correlate with parasite loads in the closely related *K. marmoratus*, see Ellison et al. 2011) and genome-wide methylation based on identification of anonymous CpG by methylation-sensitive AFLP (MS-AFLPs, previously used in non-model organisms) to identify epigenetic variation associated to parasite loads (Wenzel and Piertney 2014). Based on the Red Queen Hypothesis and previous results in *K. marmoratus*, we expected lower genetic diversity and higher parasite loads in inbred compared to outbred individuals. We also predicted higher variation in DNA methylation in relation to inbreeding and parasite loads, if methylation played an adaptive role, potentially related to immunity, in *K. hermaphroditus*.

## Methods

### Study system, field sampling and parasite screening

A total of 128 specimens of *K. hermaphroditus* were collected using hand-nets from three sampling sites on isolated mangroves on the North-eastern coast of Brazil between January and September 2015: Ceará-Mirim river – Site 1; Curimataú river – Site 2; Ipojuca river - Site 3 (Fig. 1). *K. hermaphroditus* is distributed along the Brazilian coast (Tatarenkov et al. 2017) and is typically found in shallow pools of high salinity levels (>30 ppt), clear waters and muddy substrates, where there are few other sympatric fish (Lira et al. 2015; Berbel-Filho et al. 2016). All specimens displayed the common hermaphrodite phenotype (dark colour with well-defined ocellus on the caudal fin; Costa 2011). Fish were euthanized using an overdose of tricaine methane-sulfonate (MS-222) following UK Home Office Schedule 1 (Hollands 1986), standard length was measured using a digital calliper (mm) and the whole fish were preserved in 95% ethanol at −20 for parasite screening and DNA extraction.

**Fig. 1.**
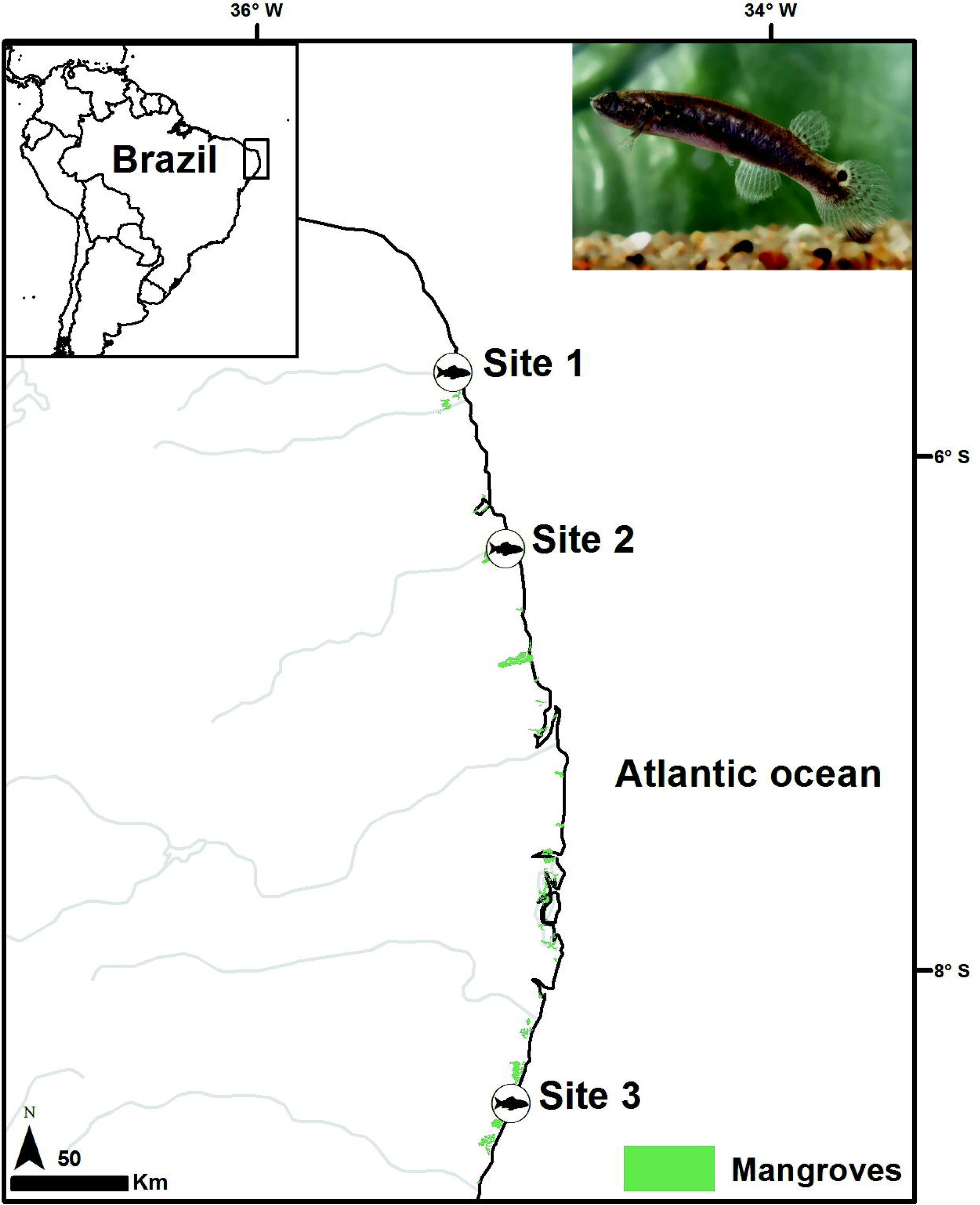
Sampling locations for *Kryptolebias hermaphroditus* (picture of a live individual on the top-right corner) in North-eastern Brazil. Ceará-Mirim river – Site 1; Curimataú river – Site 2; Ipojuca river - Site 3.

In the laboratory, fish were dissected and screened for both external and internal parasite infections using a dissecting microscope following the methods of Ellison et al. (2011). To assess the reliability of parasite screening, a subsample of five fish was examined by a different observer and the agreement was 100%. We defined parasite loads using a scaled measure of parasite abundance, where for each parasite morphotype (i), the number of parasites per individual (Ni) was divided by the maximum number found across all individuals (Nimax). The final value of the scaled parasite load represents the sum of scaled parasite loads across all parasite types. Given their uneven abundance (Table 1), this approach minimizes bias when parasite loads are heavily influenced by a very abundant parasite type (in our case bacterial cysts) (Bolnick and Stutz 2017).

**Table 1.**
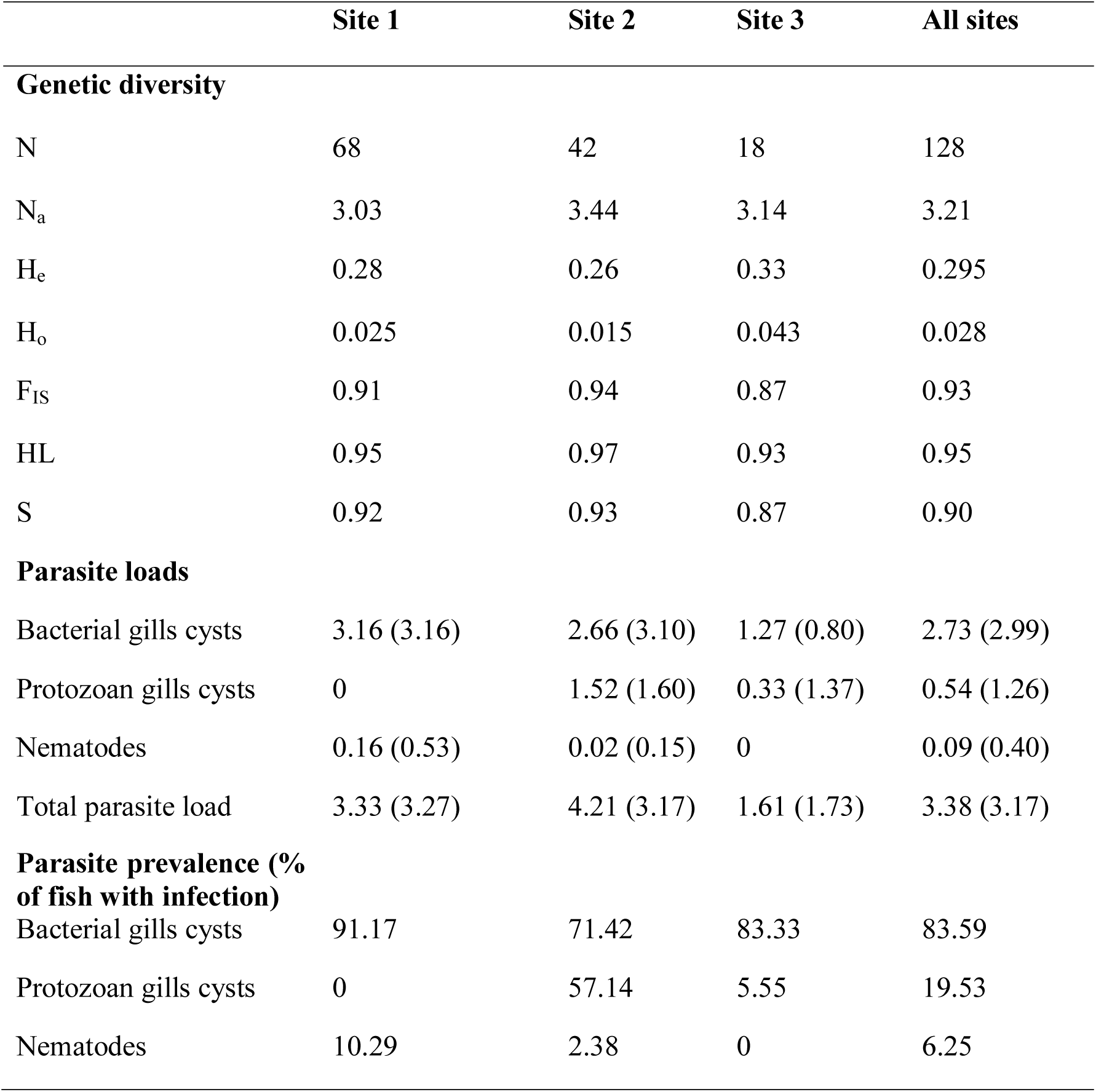
Genetic diversity (at 27 microsatellite loci), mean parasites number (standard deviation in brackets) and parasite prevalence in *Kryptolebias hermaphroditus* at sampling sites in North-eastern Brazil. N= sampling size; N_a_ = mean number of alleles of alleles; H_e_ = expected heterozygosity; H_o_ = observed heterozygosity; F_IS_ = inbreeding coefficient; HL = homozygosity by locus; S = selfing rates.

### Genetic analysis

Genomic DNA from all 128 fish was extracted from gill tissue using a Nexttec extraction kit for blood and tissue samples (Nexttec, Leverkusen, Germany). Gills are an important physical and immunological barrier to pathogens in fish (Press and Evensen, 1999), and the organ where most parasites were found (Table 1). Twenty-seven microsatellite loci (Mackiewicz et al. 2006; Tatarenkov et al. 2017) were genotyped as in Ellison et al. (2011) and screened using GeneMapper v.4.0 (Applied Biosystems, Foster City, USA). Loci were tested for linkage disequilibrium and Hardy-Weinberg equilibrium using GENEPOP v. 4.5.1 (Rousset 2008). Mean number of alleles per locus (N_ma_), observed heterozygosity (H_o_) and expected heterozygosity (H_e_) were estimated using GenALEX v. 6.5 (Peakall and Smouse 2012). The inbreeding coefficient (F_IS_) was calculated in GENEPOP. Global heterozygosity for individual fish was estimated using the homozygosity by locus index (HL) implemented in the Excel macro Cernicalin v 1.3 (Aparicio et al. 2006).

We also used the Bayesian clustering algorithm INSTRUCT (Gao et al. 2007) to estimate the optimal number of selfing lineages (k) with four simultaneous chains of 2,000,000 MCMC runs, 10 as thinning, and 100,000 of burn-in period, resulting in 100,000 interactions for each chain. The potential number of k tested ranged from 2 to 12. We used the individual *q*-values (the likelihood of membership to a particular genetic cluster or selfing lineage) from INSTRUCT to classify individuals as either selfed or outcrossed (Vähä and Primmer 2006). A threshold of *q*-value ≥0.9 was used to classify selfed individuals, while <0.9 represented hybrids between two different selfing lineages, suggesting an outcrossing event (Ellison et al. 2011; Vähä and Primmer 2006). Pairwise F_ST_ values among sampling sites and selfing lineages were estimated with Arlequin v. 3.5.2.2 (Excoffier and Lischer 2010) using 10,000 permutations. We used hierarchical analysis of molecular variance (AMOVA) to investigate population structuring among sampling sites and selfing lineages (according to individual *q*-values) using 10,000 randomizations. Differences between selfed and outcrossed groups in the total number of parasites and homozygosity by locus (microsatellites) were analysed using median Mann-Whitney rank tests in R v. 3.3.

### Epigenetic analysis

We used Methylation-Sensitive Amplified Fragment Length Polymorphisms (MS-AFLPs) to assess genome-wide DNA methylation patterns (Schrey et al. 2013). DNA extracted from gill filament tissue of 115 fish (33 classified as outcrossed and 82 as selfed according to the INSTRUCT *q*-values; 62, 36 and 17 from samplings sites 1, 2 and 3, respectively) was used for the MS-AFLPs analysis following Rodríguez López et al. (2012). A DNA aliquot of 100 ng per individual was split for digestion with two enzyme combinations: EcoRI/HpaII and EcoRI/MspI. The digested DNA was ligated to adaptors and a selective PCR was conducted using the primers ECORI-ACT: GACTGCGTACCAATTCACT and HPA-TAG: GATGAGTCTAGAACGGTAG following Ellison et al. (2015). The HpaII primer was end-labelled with 6-FAM. Fragments were run on an ABI PRISM 3100 (Applied Biosystems) and the resultant profiles were analysed using GENEMAPPER v. 4.0 (Applied Biosystems). To ensure reproducibility the following settings were applied: analysis range was 100-500 bp; minimum peak height was 100 relative fluorescence units; pass range for sizing quality: 0.75-1.0; maximum peak width: 1.5 bp. To confirm MS-AFLP reproducibility, 24 individuals (∼20% of the total; eight from each sampling site) were reanalysed and compared using the same protocols.

The R package msap v. 1. 1. 9 (Pérez-Figueroa 2013) was used to analyse MS-AFLP data. To increase reproducibility of the genotyping, we used an error threshold of 5% as suggested by Herrera and Bazaga (2010). According to the binary band patterns, each locus was classified as either methylation susceptible loci (MSL; i.e. displaying a proportion of HPA+/MSP- and/or HPA-/MSP+ sites which exceed the error threshold (5%) across all samples) or non-methylated loci (NML; if the same patterns did not exceed the error threshold) (Pérez-Figueroa 2013). MSL were used to assess epigenetic variation, while NML were used as a measure of AFLP genetic variation. Average group methylation percentages for inbreeding status were calculated using the different binary band patterns (hemimethylated pattern (HPA+/MSP-) + internal cytosine methylation pattern (HPA-/MSP+)/unmethylated pattern (HPA+/MSP+) + hypermethylation/absence of target (HPA/MSP-) *100) (Veerger et al. 2012).

Epigenetic (MSL) and genetic at AFLPs (NML) differentiation among sampling sites, selfing lineages and between outcrossed and selfed groups, was assessed by AMOVA with 10,000 randomizations. Epigenetic (MSL) and genetic (AFLP and microsatellites) differentiation among sampling sites, selfing lineages and inbreeding status was visualized by principal coordinates analysis (PCoA). Mantel tests based on distance matrices (Mantel 1967) were used to test for potential correlations between epigenetic and genetic data for MSL, NML and microsatellites using GENALEX v. 6.5 with and 10,000 permutations. To identify disproportionately differentiated methylation states, we used a F_ST_ outlier approach implemented in Bayescan 2.1 (Foll and Gaggiotti 2008; Perez-Figueroa et al. 2010), with 2×10^6^ iterations (thinning interval 20 after 20 pilot runs of 10^4^ iterations each) and a burnin of 5×10^5^. We tested for outliers based on the MSL data generated on the comparisons among sampling sites, selfing lineages and between inbreeding status (inbred or outbred).

### Statistical analyses

A Kruskal-Wallis test was used to examine the differences on scaled parasite load and bacterial cysts (the most prominent parasite) among selfing lineages. To test the relationship between genome-wide variation in DNA methylation and parasite loads, the proportion of methylated loci per individual was calculated as the proportion of loci scored as methylated over the total number of loci observed per individual (“0” for unmethylated and “1” for methylated, excluding the missing data cells per individual). We then employed a generalized linear model with a binomial link to model proportion of methylated loci as a function of scaled parasite load, selfing lineage, sampling site and inbreeding status. We repeated the analysis including only the most prominent parasite type (bacterial cysts).

Model selection was conducted using the multi-model averaging approach implemented in the R package glmulti v 1.0.7 (Calcagno and de Mazancourt 2010). We chose the minimal adequate models based on the lowest AICc values (Akaike Information Criterion corrected for small sample size), Akaike weight (W_i_) and evidence ratios (Burnham and Anderson 2004). Models (within 2 AIC units) were also reported. Predictors were checked for collinearity using *pair.panels* function in R package psych (Revelle et al. 2019). Model residuals were checked and assumptions validated.

To disentangle potential confounding effects arising from the unequal distribution of selfing lineages among sampling sites (i.e. five lineages are exclusive to a particular sampling site, Table S1), we repeated the analyses (AMOVA, Mantel test, PCoA and GLMs) for both genetic (microsatellites and AFLPs) and epigenetic (MSL) data using on individuals from Site 1 (68 individuals for microsatellites and 62 for MS-AFLPs), as this was the only site with more than two selfing lineages (Table S1).

## Results

### Parasite screening

Macroscopic parasite loads were generally low and we focused on the three most common types of parasites identified. Bacterial cysts were present on the gills and consisted of white to yellow spherical cysts circumscribed by a capsule, which resulted in hypertrophied gill filaments. They were the most common type of pathogen appearing in 83.6% of the individuals screened, with a prevalence ranging from 1 to 19 (mean=2.73, s.d.=2.99), and were more prevalent in Site 1 (mean=3.16, s.d.=3.16), followed by Site 2 (mean=2.66, s.d.=3.10) and Site 3 (mean=1.27, s.d.=0.80). The second most common macroscopic parasites were protozoan cysts, which consisted of small dark oval cysts over the gills arch and filaments. In total, 19.53% of the total number of individuals were infected with these cysts, ranging from 1 to 6 (mean=0.54, s.d.=1.26). Protozoan cysts were absent in Site 1, but present in Site 2 (mean 1.52, s.d.=1.6) and Site 3 (mean=0.33, s.d.=1.37). Finally, adult nematodes were found in the gut of only eight individuals (6.25%), ranging from 1 to 3 (mean=0.09, s.d.=0.40). Nematodes were only detected in Sites 1 (mean=0.3, s.d.=1.37) and 2 (mean=0.02, s.d.=0.15) (Fig. S1; Tables 1 and S1). Only seven individuals (5.4%) were uninfected with macroparasites. Significant differences were found on scaled parasite loads (Chi square = 32.14, p = <0.001, df = 5) and bacterial cysts (Chi square = 12.98, p = 0.01, df = 5) among selfing lineages.

### Genetic diversity and population structuring based on microsatellites

No linkage disequilibrium was detected between any pair of microsatellite loci. As expected from the high levels of self-fertilisation of the species, no loci were found to be in Hardy-Weinberg equilibrium, and all 27 microsatellite loci showed an excess of homozygotes. The global homozygosity index (HL) was very high (mean = 0.95), as well the estimated selfing rates (Table 1). At the individual level, 93 individuals (72.6%) were homozygous across all 27 microsatellite loci. However, 17 individuals (13.28%), displayed intermediate to high levels of heterozygosity (ranging from 0.13 to 0.69).

The clustering Bayesian algorithm INSTRUCT indicated that six was the most likely number of selfing lineages (k). Selfing lineage 6 was shared between two different mangroves (Site 1 with seven individuals and site 2 with one individual), separated by approximately 100 km. The other five lineages were solely represented in one of the mangroves (lineage 1 with 14 individuals, lineage 2 with 25 individuals and lineage 4 with 22 individuals in Site 1; lineage 3 with 41 individuals in Site 2; and lineage 5 with 18 individuals in Site 3) (Figs. 1 and 2; Table S1). High F_ST_ values were found both among sampling sites (mean = 0.28, s.d. = 0.02) and selfing lineages (mean=0.32, s.d. = 0.05). All pairwise comparisons were highly significant (Table S2).

**Fig. 2.**
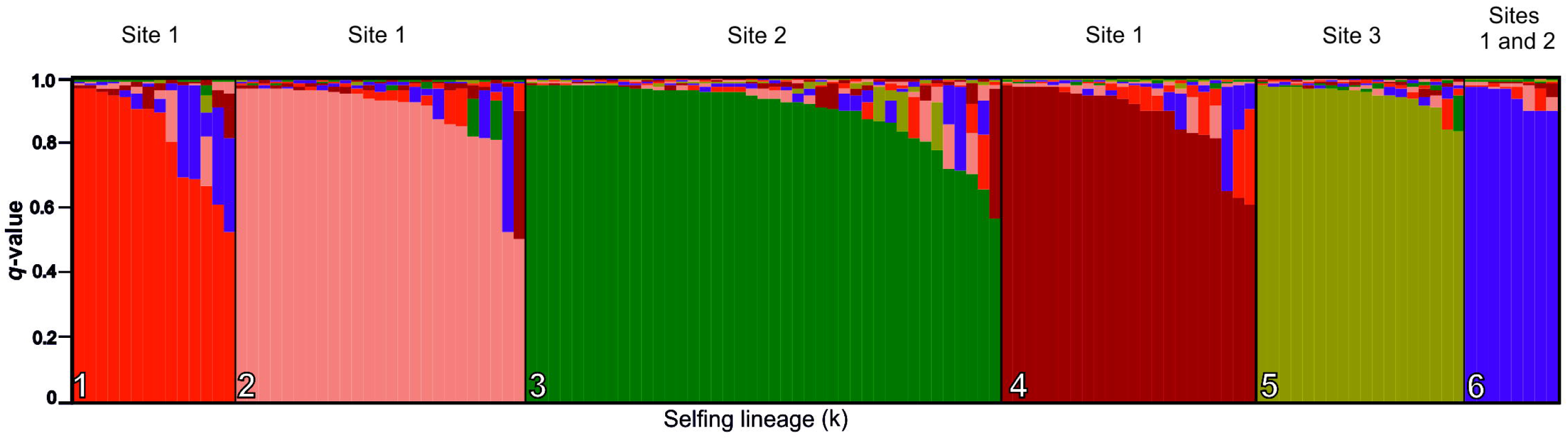
Genetic assignment of *Kryptolebias hermaphroditus* to six selfing lineages using INSTRUCT. Each individual is represented by a bar, which represents the likelihood of the individual to belong to a specific genetic cluster (colour).

Based on the *q*-values from the INSTRUCT lineages, 92 fish (71%; 46 from Site 1, 30 from Site 2, 16 from Site 3) were classified as selfed (with *q*-values ≥0.9) and 36 (29%; 22 from Site 1, 12 from Site 2 and two in Site 3) as outcrossed (with *q*-values <0.9) (Fig. 2; Table S1). Overall, outcrossed individuals had significantly lower homozygosity by locus values (at microsatellites) and total parasite loads than selfed individuals (Table 2).

**Table 2.** Homozygosity by locus (HL) (at 27 microsatellite loci), mean parasites loads (standard error in brackets) and parasite prevalence in *Kryptolebias hermaphroditus* classed as either selfed or outcrossed based on q-values from selfing lineages structure estimated using INSTRUCT. P and z-values extracted from a two median Mann-Whitney test.

|  | **Selfed** | **Outcrossed** | **z** | **P value** |
| --- | --- | --- | --- | --- |
| **Genetic diversity** |  |  |  |  |
| N | 92 | 36 |  |  |
| HL | 0.98 | 0.88 | -4.76 | **<0.001** |
| **Parasite loads** |  |  |  |  |
| Bacterial gills cysts | 3.25 (2.99) | 1.69 (2.59) |  |  |
| Protozoan gills cysts | 0.57 (1.26) | 0.47 (1.28) |  |  |
| Nematodes | 0.1 (0.4) | 0.05 (0.42) |  |  |
| Total parasite load | 3.82 (3.47) | 2.25 (1.94) | -2.84 | **0.004** |
| **Parasite prevalence (% of fish with infection)** |  |  |  |  |
| Bacterial gills cysts | 89.13 | 69.44 |  |  |
| Protozoan gills cysts | 18.47 | 22.22 |  |  |
| Nematodes | 7.6 | 2.77 |  |  |

Overall, AMOVA analyses using microsatellites indicated strong and significant differentiation among sampling sites (F_ST_ = 0.28, P = 0.001) and selfing lineages (F_ST_ = 0.32, P = 0.001), with the latter explaining more of the genetic variance than the former (Table 3). Although significant, very low genetic differentiation was found between selfed and outcrossed individuals (F_ST_ = 0.01, P = 0.002) (Table 3; Fig. S2). These patterns were also seen on PCoA analysis, with individuals generally clustering by selfing lineages in the microsatellites data (25.84% of overall variation), with individuals from lineage 4 being the most differentiated from the other lineages on Site 1. In this site, substantial overlap was found among selfing lineages and between selfed and outcrossed, despite its significant differences (F_ST_ = 0.03, P = 0.001) (Table S4; Fig. S3).

**Table 3.**
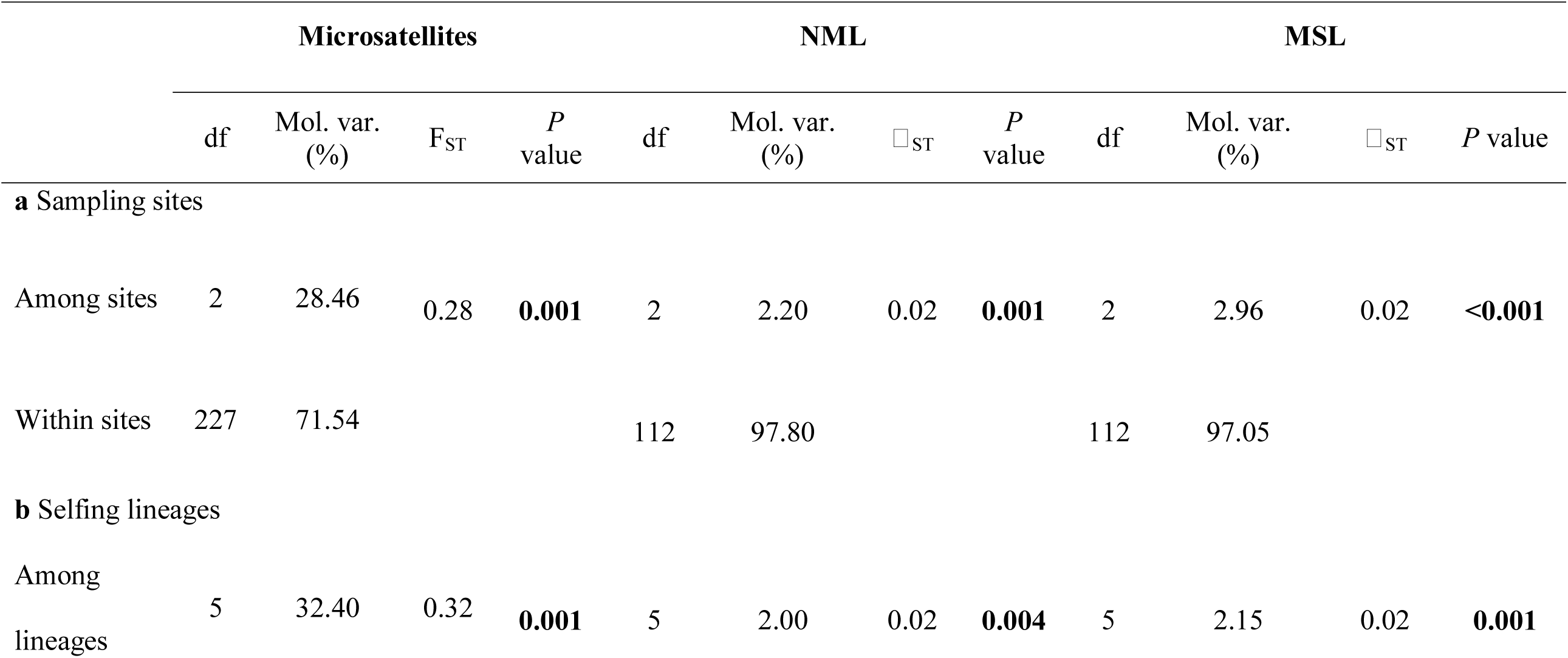
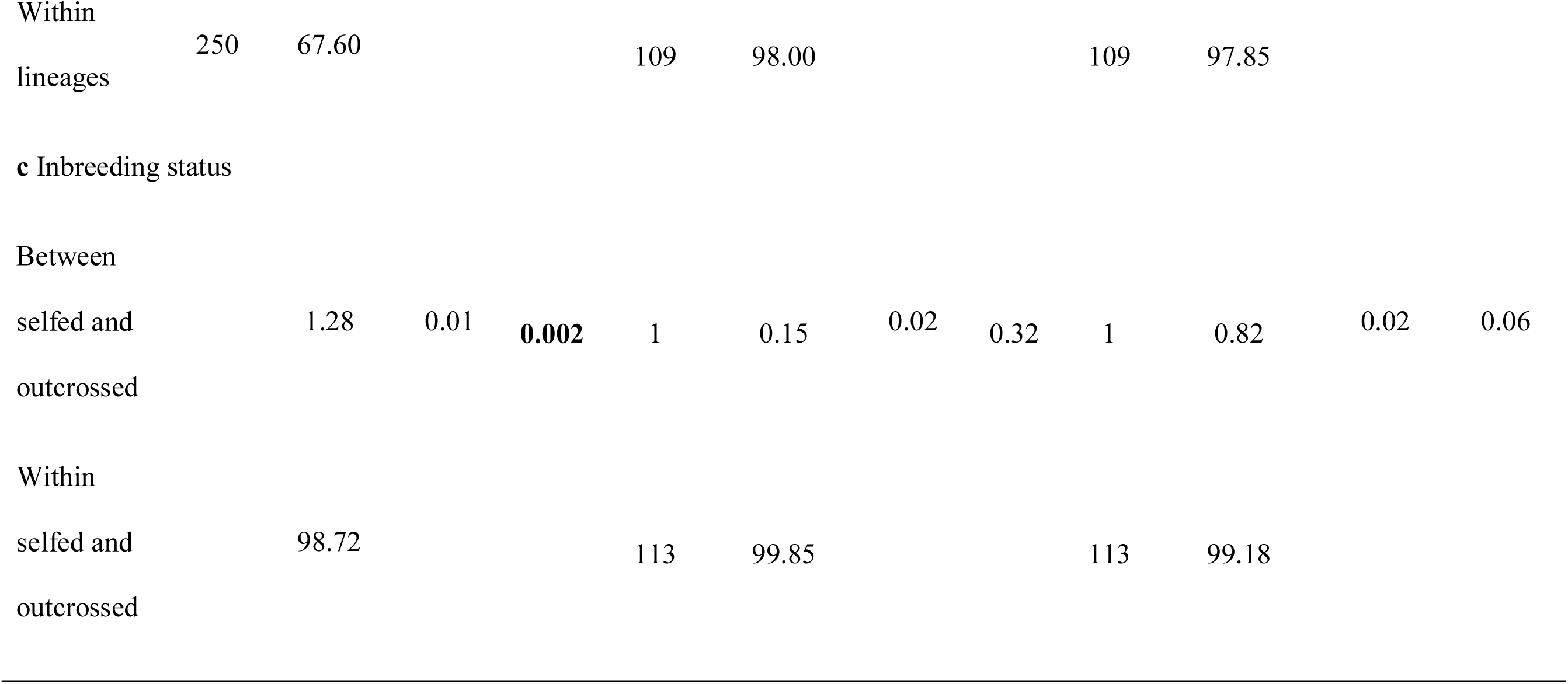
Hierarchical analysis of molecular variance (AMOVA) for microsatellites and MS-AFLPs data among **a** sampling sites **b** selfing lineages and **c** between selfed and outcrossed individuals and **b** among sampling sites in *Kryptolebias hermaphroditus*. df= degrees of freedom; SSD= sum of squared deviations; Mol. var. (%) = molecular variance percentages from variance components sigma 2; _LST_ = Phi statistics for population differentiation. *P* value derived from 10,000 permutations.

### Genetic and epigenetic variability and population structuring based on MS-AFLPs

The epigenetic analysis identified 381 MS-AFLP loci, of which 267 (70.07%) were methylation-susceptible loci (MSL) and 106 (27.82%) non-methylated loci (NML). Of the MSL loci, 236 (88.3%) were polymorphic and therefore were used for the variability analysis. Reproducibility comparisons between original and replicated genotypes for 24 individuals revealed 262 loci with an average of 10% differences (sum of number of differences between first and second set of genotypes divided by number of individuals). AMOVA analysis for reproducibility revealed no significant differences between methylation and AFLP variation patterns between original and replicated set of individuals (Table S3). Average methylation ranged from 47.51% on lineage 2 to 38.17 % on lineage 5, and was 44.82% for inbred and 45.77% for outbred individuals.

AMOVA revealed very low but significant differentiation among sampling sites, for both genetic (AFLPs: □_ST_ = 0.02, P = 0.001) and epigenetic loci (□_ST_ = 0.02, P < 0.001). Significant differentiation among selfing lineages was also found on genetic (AFLPs: □_ST_ = 0.02, P = 0.004) and epigenetic loci (□_ST_ = 0.02, P = 0.001). Overall, higher genetic and epigenetic variance was found within than between groups (Table 3). As with microsatellites, no clear genetic (at AFLPs) or epigenetic differentiation was found between selfed and outcrossed individuals (Fig. S2). There was, however, a significant positive association between epigenetic (MSL) and genetic diversity, both using AFLPs (Mantel test, r = 0.11; P = 0.002) and microsatellites (r =0.09; P= 0.001). No MSL epiloci were identified as an F_ST_ outlier in any of the comparisons.

No significant differences between selfing lineages were found among lineages for individuals from Site 1 for AFLPs genetic data (selfing lineages: □_ST=_0.008, P = 0.12) or MSL epigenetic data (selfing lineages: □_ST=_0.006, P = 0.20) (Table S4). In the PCoA, substantial overlap was found among selfing lineages and between selfed and outcrossed individuals (Fig. S3). Mantel tests between genetic and epigenetic data indicated a significant positive association between AFLPs and MSL data (r = 0.21; P <0.001), but not between microsatellites and MSL (r = −0.005; P =0.45).

### Parasite loads, genetic and epigenetic variation

According to a multi-model testing approach, the most plausible model for the proportion of methylated DNA included selfing lineage, scaled parasite load, inbreeding status and the interactions between selfing lineage and scaled parasite load and inbreeding. The proportion of methylated loci significantly varied among selfing lineages (estimate = 0.51, S. E.= 0.13, P < 0.001) and was affected by parasite loads and inbreeding status through its interactions with selfing lineage (parasite loads and selfing lineage: estimate = −0.55, S.E.=0.46, P = 0.005; inbreeding and selfing lineage interaction: estimate = −1.64, S.E.=0.14, P = 0.04) (Fig 3b-c; Tables 4 and S7). The second most likely model (ΔAICc=1.00) included only selfing lineage (estimate = −0.43, S. E.=0.08, P < 0.001) and the interactions between inbreeding and selfing lineage (estimate = −1.10, S. E.=0.12, P = 0.04) as significant predictors. However, this model explained substantially less of the overall variation compared to the first model (weight: 0.17 vs. 0.28). and was 1.39 times less likely than the first one (Table S5).

**Fig. 3.**
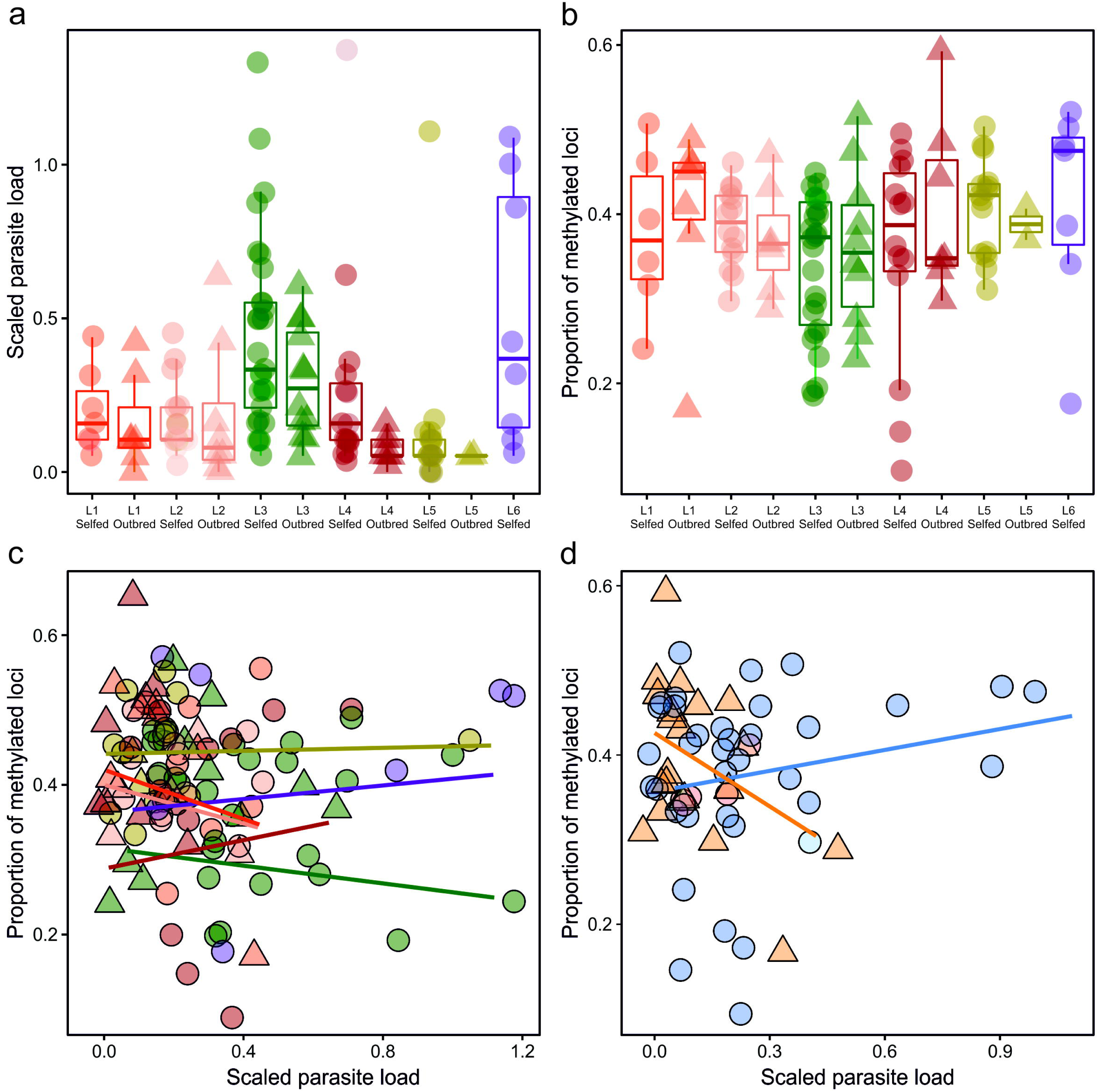
Relationships between **a** scaled parasite load across selfing lineages and inbreeding status **b** proportion of methylated loci across selfing lineage and inbreeding status (selfed or outcrossed) **c** Proportion of methylated loci and selfing lineages and scaled parasite loads **d** proportion of methylated loci across inbreeding status for sampling site 1 individuals. Circles for selfed, triangles for outcrossed individuals. Red = selfing lineage 1 (site 1); salmon = selfing lineage 2 (site 1); green = selfing lineage 3 (site 2); brown = selfing lineage 4 (site 1); yellow = selfing lineage 5 (site 3); purple = selfing lineage 6 (sites 1 and 2); orange = outcrossed individuals; blue = selfed individuals.

**Table 4.**
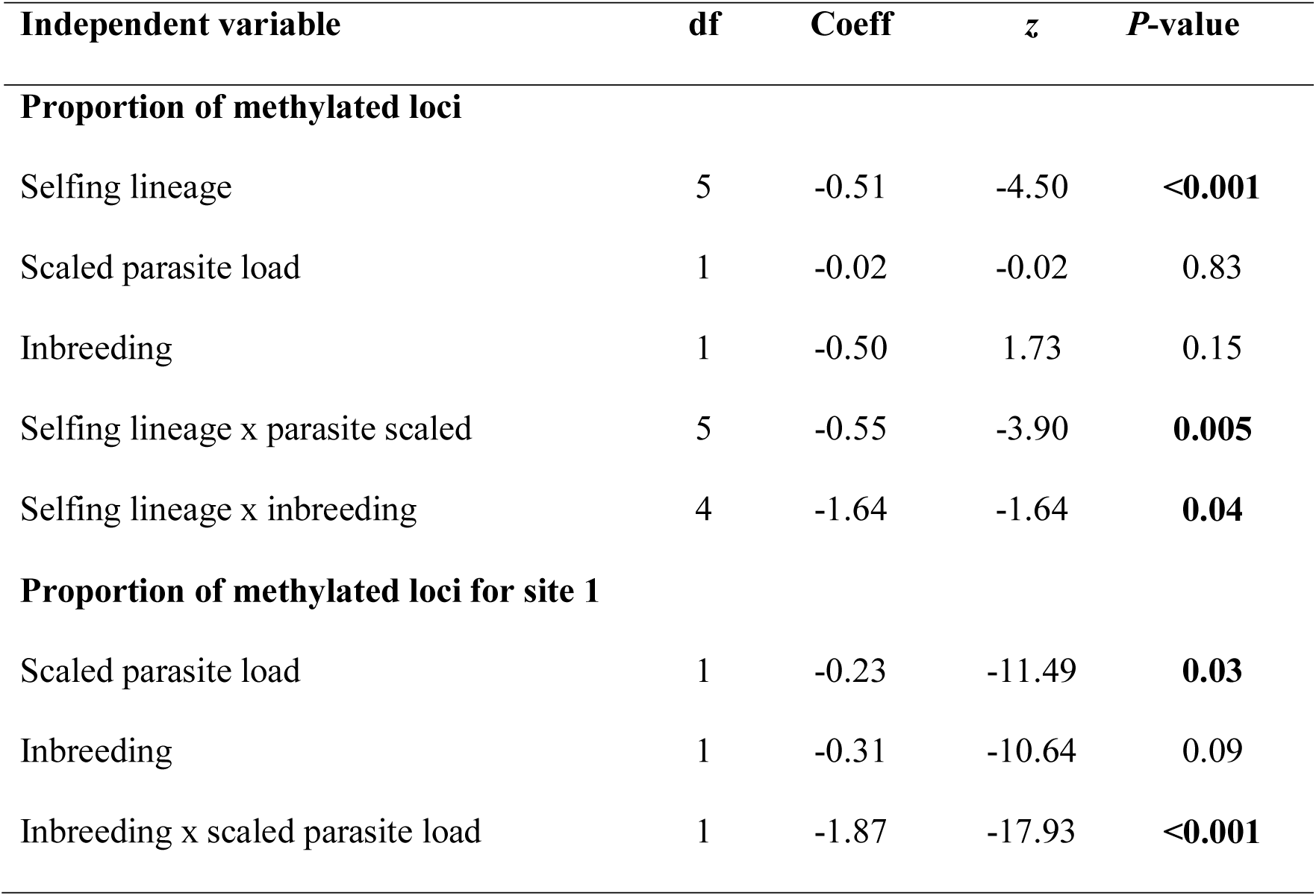
Results of the best-fitting generalized linear models for proportion of methylated loci (binomial distribution) in *Kryptolebias hermaphroditus*, using the multi-model averaging approach (see appendix for the full model comparisons). df= degrees of freedom; Coeff = mean coefficient estimates.

Overall, the results of the single-taxa models (using number bacterial cysts) were very similar to those for scaled parasite loads. The best model to explain the proportion of methylated loci included selfing lineage, and the interactions between selfing lineage and bacterial cysts, and selfing lineage and inbreeding (Table S6).

When using only individuals from Site 1 (to remove any potential confounding effect between sampling site and selfing lineages) for the proportion of methylated loci the model with the lowest AIC indicated that selfing lineage, inbreeding and the interactions between inbreeding and selfing lineage and inbreeding and scaled parasite loads were all significant predictors (Table S7). However, the second best-fitting model (ΔAICc = 0.02) explained the same amount of variation (weight=0.39) and the evidence ratio (−0.66) suggested that it was more likely (evidence ratio of 1.50) than the first model. This second model indicated that overall, the proportion of methylated DNA significantly increased with scaled parasite loads (estimate = 0.43, S. E.= 0.11, P = 0.03) and that DNA methylation levels were also affected by the interaction between scaled parasite loads and inbreeding (estimate = −1.29, S. E.=0.38, P <0.001), with inbred individuals having increased methylation levels with increased parasite loads, while outbred individuals had decreased methylation levels with increased parasite loads (Fig. 3d; Table 4).

## Discussion

Overall, our results did not indicate significant differences in genome-wide DNA methylation variation between selfed and outcrossed individuals, and our models only identified inbreeding significantly related to DNA methylation via its interaction with selfing lineage (all sampling sites) and parasites (at the local scale in Site 1). Higher variation in DNA methylation has been reported for clonal and inbred individuals (Nakamura and Hosaka 2010; Massicotte and Angers 2012; Richards et al. 2012; Liebl et al. 2013; Veerger et al. 2012), and has been interpreted as an adaptive mechanism to compensate for low genetic variarion (Schrey et al. 2012), or as a potential consequence of inbreeding (as in Vergeer et al. 2012) responsible, at least in part, for inbreeding depression (Nakamura and Hosaka 2010; Vergeer et al. 2012). Yet, our results suggest that, at least in this species, either inbreeding does not affect genome-wide DNA methylation variation or it does in a gene-specific manner (Venney et al. 2016), although further research would be needed to address this question.

We found that that the different selfing lineages of *Kryptolebias hermaphroditus* distributed in three sampling sites of north-eastern Brazil differed significantly in parasite loads and genetic composition, which might indicate specific interactions between host genotypes and parasites (Dybdhal and Lively 1998; Ebert 2008). Previous studies on mangrove killifishes had identified extensive genetic structuring both between (Tatarenkov et al. 2015; 2017) and within mangrove systems even at close geographical proximity (Tatarenkov et al. 2007; 2012; Ellison et al. 2012), as a consequence of the self-fertilising nature of these fish. We found strong evidence of genetic structuring between sampling sites and selfing lineages using microsatellites, but lower differentiation for AFLP genetic markers (likely due to the different mutation rate of the markers) and epigenetic markers (MS- AFLPs). Overall, inbred individuals (with lower heterozygosity) harboured higher parasite loads than their outcrossed counterparts, supporting the prediction that low genetic diversity due to self-fertilisation may reduce fitness (considering parasite loads as a proxy for pathogen pressure), as for other mixed-mating species (King et al. 2011; Ellison et al. 2011; Lively and Morran 2014). Extensive periods of self-fertilisation can reduce offspring fitness due to the accumulation of deleterious alleles and inbreeding depression (Charlesworth et al. 1993). Species with mixed-mating seem to overcome these problems through occasional outcrossing (Ellison et al. 2011; Morran et al. 2011), which can generate genetic diversity to face natural enemies, such as parasites (Lively 2014). Here, the relationship between parasites and inbreeding status (selfed or outcrossed) suggests that outcrossing might confer a fitness advantage (in terms of parasite loads), even when it occurs at very low frequencies (Ellison et al. 2011). However, despite the adaptive potential of outcrossing, the main reproductive strategy of *K*. *hermaphroditus* seems to be self-fertilisation (Tatarenkov et al. 2017). This suggests that other evolutionary mechanisms may be balancing the harmful effects of parasite infections, or that parasite selection is of low (Lively 2014), as theory predicts that low selection levels imposed by natural enemies are consistent with the maintenance of asexual reproduction (Ladle et al. 1993; Judson 1997). For example, in the mixed-mating *Potamopyrgus* snails, the oldest asexual lineages are restricted to populations where parasites are rare (Neiman et al. 2005). Thus, the low number of parasites found in *K*. *hermaphroditus* (i.e. mean of 3.38 parasites per individual compared to 22.41 of *K. marmoratus* in Belize; Ellison et al. 2011), may explain the high prevalence of selfing in *K*. *hermaphroditus*. The long-term persistence of self-fertilising organisms suggests that non-genetic mechanisms may play a role in generating adaptive responses to environmental change and compensate for low genetic variation (Shrey et al. 2012; Liebl et al. 2013; Douhovnikoff and Dodd 2015; Hu et al. 2018). Using data from all sampling sites, we found that genome-wide DNA methylation was strongly influenced by selfing lineage and only at a smaller scale by inbreeding through its interaction with selfing lineage (Bell et al. 2011; Bjornsson et al. 2008; Gertz et al. 2011; Dubin et al. 2015). Strong epigenetic differences between selfing lines had been identified previously in *K. marmoratus* (see Ellison et al. 2015), indicating an important role of the genetic background in the epigenetic variation of these species. In addition, we also found a significant correlation between DNA methylation and genetic variation (at both AFLP and microsatellites data), suggesting that autonomous variation in DNA methylation may be limited in this study system (Dubin et al. 2015).

DNA methylation can interact with genotypes in a genotype-by-environment manner to generate plastic responses (Herman and Sultan 2016). Several abiotic and biotic factors, including parasites (Norouzitallab et al. 2014; Hu et al. 2018), are known to influence DNA methylation, however information on how DNA methylation varies across different genetic backgrounds is still scarce. Our results showed that genome-wide DNA methylation levels for all sampling sites were significantly influenced by parasite loads through the interaction with selfing lineage, suggesting a potential genotype-by-environment interaction on parasites responses. Yet, as most of the selfing lineages were exclusive to specific sampling sites, we could completely discard confounding effects between both variables. In fact, selfing lineage did not affect genome-wide DNA methylation levels in Site 1, but only parasites and their interaction with inbreeding status. The anonymous nature of our genetic and epigenetic markers is a limiting factor to infer the potential adaptive/functional role of the DNA methylation variation in response to parasites. Further analyses, ideally under controlled experimental conditions and using higher resolution sequencing methods (i.e. whole-genome bisulfite sequencing, RNAseq), should help to clarify how reduced DNA methylation may affect immune responses in mixed-mating *Kryptolebias* species.

The relationship between parasite loads and outcrossing seems to be common to several mixed-mating species (Steets et al. 2007; Ellison et al. 2011; King et al. 2011) in addition to *K. hermaphroditus*, suggesting that the influence of parasites in the regulation of mixed-mating could be generalised. The extent of this relationship, however, may depend on the severity of the selection imposed by coevolving parasites (Lively and Morran 2014). Our results indicate that genotype composition (and its interaction with inbreeding) may be important in DNA methylation responses to environmental variation in wild populations, and that, if DNA methylation responded in a genotypic-specific manner to parasites pressures, it could contribute to local adaptation (Foust et al. 2016; Smith et al. 2016). The mangrove killifish, with its naturally inbred populations and marked methylation differences between populations and genotypes, represents an ideal model to analyse the relative roles of genetic and epigenetic diversity in modulating local adaptation.

## Acknowledgements

This work was supported by the Conselho Nacional de Desenvolvimento Científico e Tecnológico (CNPq) through a Science without Borders scholarship (233161/2014-7). This work was also funded by the National Geographic/Waitt foundation grant (W461-16). We thank Amy Ellison for the help with parasite screening. We are thankful for the LISE laboratory team and CTA facilities from Universidade Federal do Rio Grande do Norte for support during fieldwork.

## Compliance with ethical standards

All the experiments in this study have been conducted following Home Office regulations, approved by Swansea, Cardiff and UFRN (CEUA) Universities Animal Ethics Committees, and under sampling permit number 30532-1/2011 issued by ICMBio/SISBIO. The authors declare they have no conflict of interest.

## Data availability

Data available from Dryad Digital Repository: https://doi.org/XXX/XXXX

## Authors contributions

SC, WMB-F & CGL conceived the work; SMQL planned the field work and conducted the sampling together with WMB-F, CGL & SC; WMB-F did the microsatellite and parasite screening, with contributions from JC; WMB-F and PM performed the MS-AFLP analyses. WMB-F analysed the data with the contribution of SC, CGL and PM. WMB-F and SC wrote the paper with contributions from all authors.

